# Recovery from Spreading Depolarization is slowed by aging and accelerated by antioxidant treatment in locusts

**DOI:** 10.1101/2024.10.10.617596

**Authors:** R. Meldrum Robertson, Yuyang Wang

**Author notes:** Address: 3118 Biosciences Complex, Queen’s University, Kingston, Ontario, K7L 3N6 Canada.

## Abstract

Spreading depolarization (SD) temporarily shuts down neural processing in nervous systems with effective blood brain barriers. In mammals this is usually pathological in response to energetic stress. In insects a very similar process is induced by abiotic environmental stressors and can be beneficial by conserving energy. Age is a critical factor for predicting the consequences of SD in humans. We investigated the effect of aging on SD in an insect model of SD and explored the contribution of oxidative stress. Aging slowed the recovery of intact locusts from asphyxia by water submersion. In semi-intact preparations we monitored SD by recording the DC potential across the blood brain barrier in response to bath application of the Na^+^/K^+^-ATPase inhibitor, ouabain. Treatment with ouabain induced changes to the DC potential that could be separated into two distinct components: a slow, permanent negative shift, similar to the negative ultraslow potential recorded in mammals and human patients, as well as rapid, reversible negative DC shifts (SD events). Aging had no effect on the slow shift but increased the duration of SD events from ∼0.6 minutes in young locusts to ∼0.9 minutes in old ones. This was accompanied by a decrease in the rate of recovery of DC potential at the end of the SD event, from ∼1.5 mV/s (young) to ∼0.6 mV/s (old). An attempt to generate oxidative stress using rotenone was unsuccessful, but pretreatment with the antioxidant, N-acetylcysteine amide, had opposite effects to those of aging, reducing duration (control ∼1.1 minutes, NACA ∼0.7 minutes) and increasing rate of recovery (control ∼0.5 mV/s, NACA ∼1.0 mV/s) suggesting that it prevented oxidative damage occurring during the ouabain treatment. The antioxidant also reduced the rate of the slow negative shift. We propose that the aging locust nervous system is more vulnerable to stress due to a prior accumulation of oxidative damage. Our findings also strengthen the notion that insects provide useful models for the investigation of cellular and molecular mechanisms underlying SD in mammals.

**Significance Statement:** Anoxia and similar energetic crises trigger a shutdown of central neural processing in a process of spreading depolarization (SD) which is generally pathological in mammals and protective in insects. We show that some variability in the consequences of SD in an insect model can be attributed to age, such that older animals are slower to recover. Moreover, preventing oxidative stress with an antioxidant speeds recovery. These findings demonstrate a role for oxidative stress in contributing to the vulnerability of the aging insect CNS in energetic emergencies.

Graphical Abstract of Robertson and Wang - Locust Spreading Depolarization

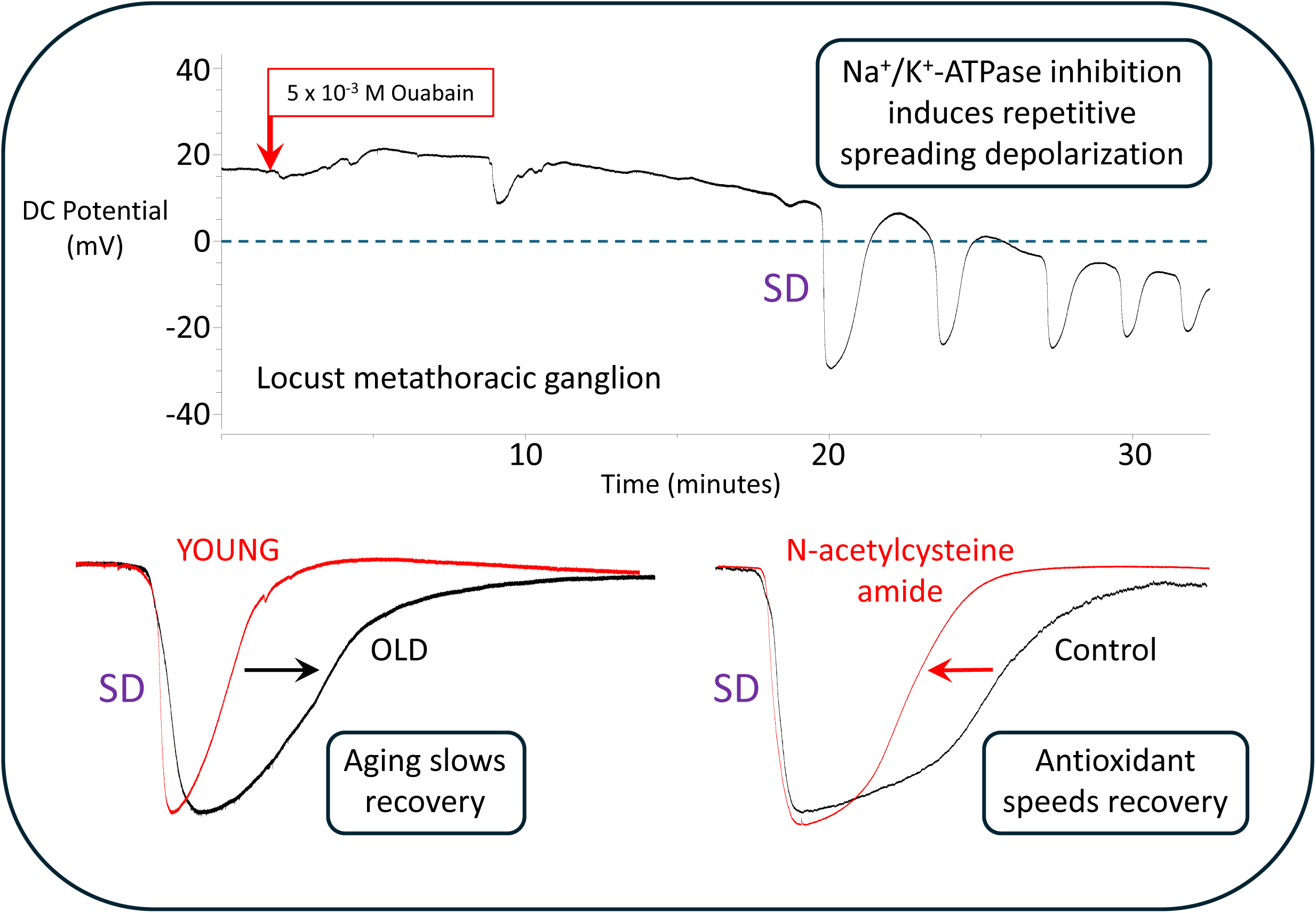

## Introduction

Complex behaviour depends on rapid and precise neural processing that, in turn, depends on the generation of electrical signals and maintenance of ion concentration gradients. The homeostatic mechanisms underlying this are energetically expensive and vulnerable to failure under metabolic stress. Very often, in a wide variety of animals, such failure manifests as a process of spreading depolarization (SD) of neural cells as ion gradients collapse. In mammals, including humans, the consequences are generally pathological, resulting in a continuum of negative outcomes from migraine aura to neuron death following traumatic brain injury (Dreier and Reiffurth, 2015; Andrew et al., 2022). In insects, however, the consequences are more benign, and SD can be considered as a means of temporary energy conservation during environmental challenges (Robertson et al., 2020; Robertson et al., 2023). Nevertheless, much evidence points to the similarity of the underlying mechanisms in mammals and insects (Rodgers et al., 2010; Spong et al., 2016b), Indeed, recent research in both mammals and insects suggests that the SD channel may in fact be the Na^+^-K^+^-ATPase (NKA) converted from a pump to a channel (Kim et al., 2024; Wang et al., 2024), a property of NKA that is well established (Gadsby et al., 2009; Rossini and Bigiani, 2011). Given the importance of age for pathological outcomes in humans, we were interested in aging as a source of variability in the progression of SD in the locust model.

Aging refers to the gradual deterioration of function over time because of the accumulation of molecular and cellular damage. Amongst the many factors contributing to an aging phenotype, oxidative damage due to reactive oxygen species (ROS) plays a major role (Finkel and Holbrook, 2000; Li et al., 2024). The aging mammalian brain is <u>less</u> susceptible to SD, but each SD event tends to be prolonged, suggesting a depletion of resources required for recovery (Clark et al., 2014; Menyhárt et al., 2015; Guedes and Abadie-Guedes, 2019; Hertelendy et al., 2019). This may result from the age-related decline in brain NKA activity as a consequence of enhanced oxidative damage (Chakraborty et al., 2003). SD causes increased levels of ROS in neural tissue (Viggiano et al., 2011; Aboghazleh et al., 2021) and ROS can trigger SD (Malkov et al., 2014) and increase its speed of propagation (Germano et al., 2022). Such oxidative stress can induce long-term mitochondrial injury and breakdown of the blood brain barrier (van Hameren et al., 2024). In *Drosophila*, on the other hand, the aged brain is <u>more</u> susceptible to SD induced by application of the NKA inhibitor, ouabain, evidenced primarily by a decrease in the latency to an increased number of SD events, though these were similarly of longer duration (Spong et al., 2016a). Also, old flies take longer than young flies to recover from submersion or nitrogen anoxia (Benasayag-Meszaros et al., 2015), a phenotype that can be genetically manipulated by reducing antioxidant capacity (Suthakaran et al., 2021).

SD can be monitored with a wide variety of different phenomena bespeaking the profound influence it has on neural function: depression of neural activity, cell swelling, extracellular ion concentrations (primarily potassium, [K^+^]_o_), extracellular transmitter concentrations, intracellular membrane potentials, extracellular DC potentials etc. A relatively simple technique in insects is to record the extracellular potential developed across the sheath of a ganglion (Schofield and Treherne, 1984), which incorporates a mechanical barrier for structural support (the lamella) and a barrier to ion diffusion (the perineurial and subperineurial glial layers) (Limmer et al., 2014). Such recordings of the transperineurial potential (TPP) can be variable and difficult to interpret because of the interplay of mechanisms that contribute to the TPP (e.g. NKA pump current, proton pump current, ion concentrations, ion conductances etc.) (Robertson and Van Dusen, 2021). However, deciphering the trajectory of a TPP trace during the induction of SD may help to disentangle the relative roles of different contributory processes and generate hypotheses that can be tested with more precision in other, molecular genetic, models (e.g. *Drosophila* (Spong et al., 2016a; Spong et al., 2017; Suthakaran et al., 2021)).

In locusts, bath application of ouabain generates SD evident as repetitive surges of extracellular potassium [K^+^]o (Rodgers et al., 2007) and as negative shifts of TPP (Spong et al., 2016c). More recently it has been established that ouabain treatment also causes a slow negative shift of TPP (reversal) from ∼ +13 mV to stabilize at ∼ −31 mV (Van Dusen et al., 2020). This reduces the amplitude of concurrent anoxia-induced SDs by changing the background TPP, thus rendering SD amplitude of little use in characterizing the severity of ouabain-induced SD. It is unclear how this TPP reversal is associated with SD.

In the current study we tested the hypothesis that aging, via oxidative stress, promotes vulnerability to anoxia and SD in locusts. We used submersion anoxia to examine the effect of aging on anoxia tolerance (resistance and recovery) of intact *Locusta migratoria*. Using bath application of ouabain to generate repetitive SD in semi-intact preparations, we analyzed TPP recordings in different preparations and simultaneous TPP recordings from right and left sides of the same ganglion to determine if TPP reversal and SD are generated by the same mechanism. In addition, we investigated the effect of aging on ouabain-induced TPP reversal and parameters of SD events. Finally, we manipulated oxidative stress using rotenone to inhibit mitochondrial complex I and generate ROS (Coulom and Birman, 2004; Radad et al., 2019; Gu and Chen, 2020) and N-acetylcysteine amide (NACA) as a potent antioxidant that crosses the blood brain barrier (Amer et al., 2008; Liu et al., 2017; Chung et al., 2020).

## Material and Methods

### Animals

Migratory locusts, *Locusta migratoria*, were obtained from a long-established, breeding colony maintained in the Department of Biology at Queen’s University. They were kept in crowded cages to ensure they remained in the gregarious phase. Incandescent 40 watt light bulbs in the cages on a 12:12 light:dark cycle heated the cages to ∼30 °C during the day, returning to room temperature (∼26 °C) at night. Locusts were fed daily on fresh wheat grass, supplemented *ad libitum* with a dry mixture of bran, yeast, and milk powder. Adult cages were marked with the day’s date when the first animals moulted from 5^th^ instar nymphs to winged adults, and fresh cages were established weekly. Hence, on the day of an experiment, a locust’s age could be roughly determined as number of days post-imaginal moult and this would be a maximum for locusts from any particular cage (i.e. in any cage the first animals to moult were the oldest and determined the age for the cohort of locusts in that cage). In our colony, adult maturation takes 10 to 14 days from the final moult until animals start to reproduce and the CNS is fully developed (Gee and Robertson, 1994; Gray and Robertson, 1994; Gray and Robertson, 1996). Hence, for the current experiments, young mature adults were 3 to 4 weeks and old adults were > 7 weeks past the imaginal moult. Locusts were weighed before each experiment. To control for changes in colony conditions and time of day, control and experimental locusts from the same cage were used on the same day and the order of experiments was swapped on different days.

### Whole Animal Submersion

For these experiments only males were used to minimize variability due to different sizes (mean ± SD for mature adults in the colony: males – 1.3 ± 0.2 g; females – 2.0 ± 0.3 g). Also, we have already shown that males enter a submersion coma sooner than females and recover more slowly (Hou et al., 2014) precluding a need to establish a sex difference. Locusts were placed in separate compartments of a perforated plastic container and submerged in room temperature water. Their movements were visually monitored, and the time to an anoxic coma (“Time to coma”) was measured as the point at which the animal finished convulsing and remained motionless. All animals were kept underwater for 30 minutes, measured from the initial time of submersion, and then removed, dried, and placed on an absorbent paper towel at room temperature. Behaviour was again visually monitored for a return of regular abdominal pumping (“Time to ventilate”) and a full regaining of nervous function indicated by an abrupt “flipping over” of the animal into a standing position (“Time to stand”) (Hou et al., 2014).

### Electrophysiological Recording

We used a semi-intact preparation (Robertson and Pearson, 1982) to expose the thoracic ganglia. The wings, legs and pronotum were cut off and the locust was pinned to a cork board. The thorax and abdomen were spread open after a dorsal midline incision. The salivary glands and gonads were removed. The gut was either cut posteriorly and lifted out of the body cavity or, in later experiments, left intact and pulled to one side out of the thorax. The ventral diaphragm and other muscles overlying the ganglia were removed. Nerves 3, 4 and 5 on both sides of the metathoracic ganglion were cut close to the ganglion to improve access of pharmacological agents into the neuropil. The preparation was bathed in standard locust saline (in mM: 147 NaCl, 10 KCl, 4 CaCl_2_, 3 NaOH, 10 HEPES buffer; pH adjusted to 7.2; chemicals from Sigma-Aldrich) and grounded with a chlorided silver wire in the abdomen. For nitrogen anoxia the preparation was placed in a chamber that could be filled with nitrogen gas (Robertson and Van Dusen, 2021).

Extracellular potassium concentration in the neuropil ([K^+^]_o_) was recorded using K^+^-sensitive glass electrodes pulled from silanized glass capillary tubes. Tips were filled with Potassium Iononphore I-Cocktail B (5% valinomycin; Sigma-Aldrich) and shafts were back-filled with 500 mM KCl (for further details see (Rodgers et al., 2007)). The extracellular reference electrodes were pulled from glass capillaries to a resistance of 5-7 Mohm, back-filled with 3 M KCl. K^+^ electrodes were calibrated using 15 mM and 150 mM KCl before use. Signals from the K^+^ electrode were amplified using a DUO773 amplifier (WPI Inc.; Sarasota). Recording of the transperineurial potential (TPP) was with extracellular glass electrodes (prepared as above) and signals were amplified with a model 1600 Neuroprobe amplifier (AM-Systems). In some experiments, ventilatory motor activity was recorded with a glass suction electrode on the median nerve with signals amplified by a model 1700 differential AC amplifier (AM Systems). All electrical signals were digitized (1440A digitizer, Molecular Devices) with a sampling rate of 100 kHz and recorded (Axoscope 10.7) for later analysis.

### Pharmacology

Pharmacological agents were purchased from Sigma-Aldrich. We used a stock solution of 10^-2^ M ouabain in locust saline to inhibit NKA and generate repetitive SD (Van Dusen et al., 2020; Robertson and Van Dusen, 2021). The membrane-permeable antioxidant, NACA (Chung et al., 2020) was stored frozen in aliquots prior to use and used at 2 x 10^-3^ M in locust saline. Rotenone was dissolved in 1 mL of DMSO and diluted with 100 mL locust saline prior to freezing in aliquots, and was used at 0.5 x 10^-3^ M and 1 x 10^-3^ M. Monosodium iodoacetate (MIA), an inhibitor of glycolysis, was similarly frozen in aliquots and used at 5 x 10^-3^ M (Wang 王宇扬 et al., 2023).

Pharmacological pretreatment of the thoracic ganglia followed a standard procedure. After the initial dissection, the saline was removed from the thoracic cavity, which was dried with a paper tissue. Then 0.5 mL of pretreatment solution (saline, NACA, rotenone) was added and nerve roots 3, 4, and 5 of the metathoracic ganglion were cut to improve access to the neuropil. After 30 minutes, the extracellular glass electrode was inserted into the metathoracic ganglion, approximately in the centre of one side of the metathoracic neuromere. Finally, 0.5 mL of 10^-2^ M ouabain was added to the bathing solution, resulting in a ouabain concentration of 5 x 10^-3^ M, and the TPP was recorded for 30 minutes.

### Data analysis

We used Clampfit 10.7 (Molecular Devices) to analyze our recordings and Sigmaplot 15 (Systat, Grafiti LLC) or GraphPad Prism 10 (GraphPad Software, Boston, MA) to perform statistical tests and generate graphs. Any voltage drift of the recording during the experiment was corrected prior to analysis. Application of ouabain to the CNS generates repetitive SD that can be recorded as abrupt, reversible increases in [K^+^]_o_ (Rodgers et al., 2007) or abrupt, reversible negative shifts of TPP (Van Dusen et al., 2020). To characterize ouabain-induced changes in TPP we made the following measurements: initial TPP at penetration; background TPP (i.e. excluding abrupt SD events) after 30 minutes of ouabain treatment (TPP30); time to the first SD; and total number of SDs in 30 minutes. For the first SD we measured: amplitude; slope of the TPP trajectory at the beginning of an event (slope on); duration at half-amplitude of the event; slope of the TPP trajectory at the end of an event (slope off); and the time constant of an exponential fit to the TPP recovery trajectory (tau). Outliers were detected using Grubbs test (GraphPad) online and removed prior to further analysis. Data were tested for normality (Shapiro-Wilk test) and equal variance (Brown-Forsythe test) prior to statistical comparison using standard tests as appropriate and reported in the Results text. Averages are reported as mean ± standard deviation (SD) for descriptive data and as mean ± standard error (SE) for statistical comparisons. Graphs show scatter plots of correlations or box plots indicating the median and 25^th^ and 75^th^ percentiles with whiskers to the minimum and maximum values and overlaid with the individual data points.

## Results

### Water submersion

Ten young and ten old locusts were tested for their ability to withstand and recover from an anoxic coma induced by a 30-minute submersion in water. One old locust was removed as an outlier. The measurements were differentially affected by mass (**Fig. 1**). In young animals, mass was negatively correlated with time to coma (Pearson r = −0.857, P = 0.0016) but not the recovery measures. In old animals, mass was negatively correlated with recovery measures (time to ventilate: Pearson r = −0.85, P = 0.0037; time to stand: Pearson r = −0.80, P = 0.009) but not time to coma. Age and mass both affected time to coma (**Fig. 1A**; One Way ANCOVA: P_age_ = 0.01; P_mass_ = 0.002; P_age x mass_ = 0.007), time to ventilate (**Fig. 1B**; One Way ANCOVA: P_age_ < 0.001; P_mass_ = 0.013; P_age x mass_ < 0.001), and time to stand (**Fig. 1C**; One Way ANCOVA: P_age_ < 0.001; P_mass_ = 0.007; P_age x mass_ = 0.005).

**Figure 1:**
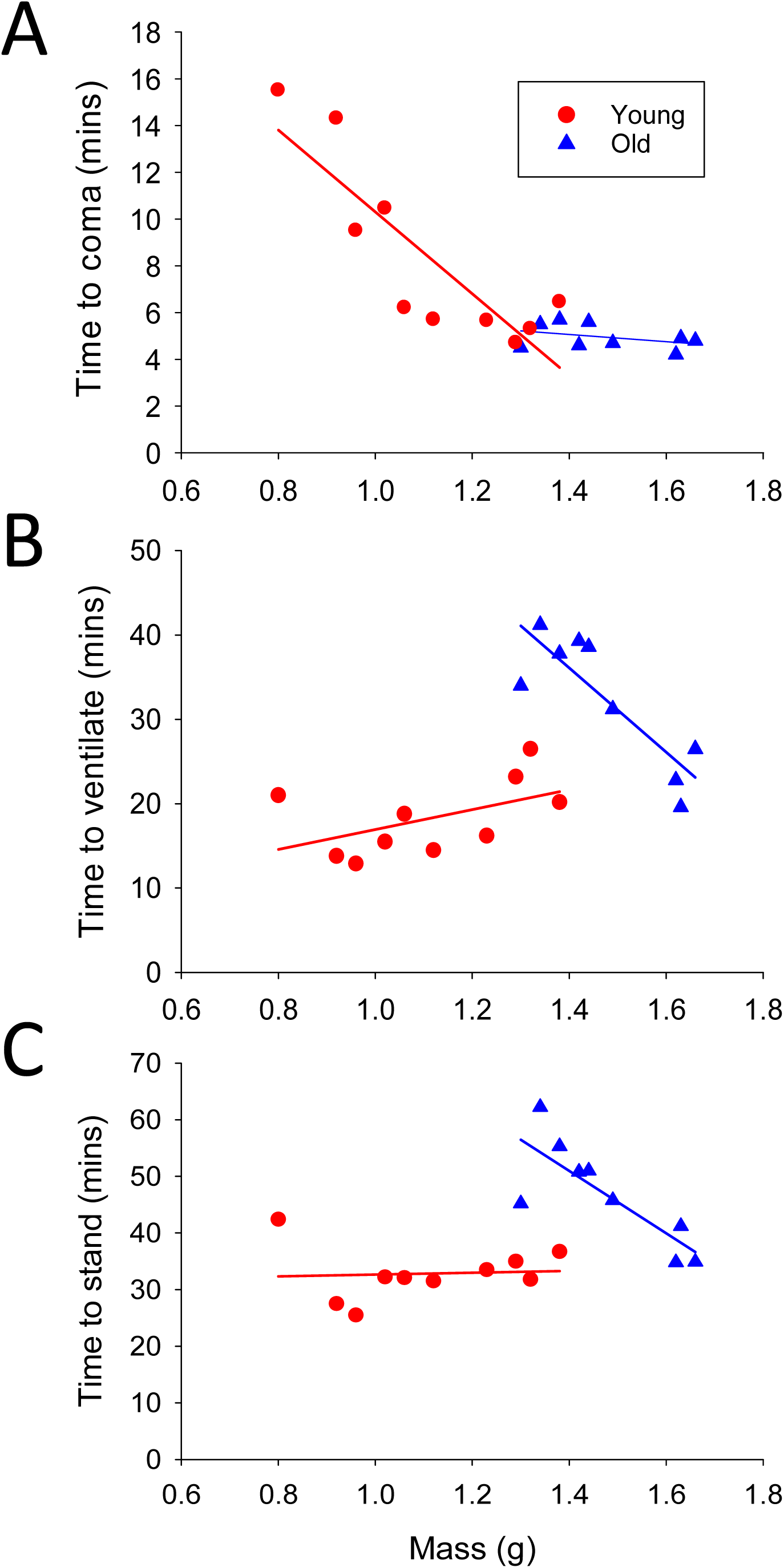
Aging slowed recovery from anoxic coma. **A.** Young and old male locusts of approximately equal mass (1.0-1.5 g) entered anoxic coma around the same time after water submersion. **B.** Old locusts took longer to recover ventilatory movements after removal from the water than young locusts of approximately equal mass. **C.** Old locusts took longer to recover the ability to stand after removal from the water than young locusts of approximately equal mass. Young – red circles; Old – blue triangles. Statistical comparisons in text.

### Ouabain SD phenomenology

Ouabain treatment of a thoracic ganglion with nerve roots cut usually induced repetitive SD superimposed on a gradual reversal of the TPP from positive to negative (**Fig. 2Ai**). Frequently in control animals repetitive SD occurred without TPP reversal (**Fig. 2Aii**). Rarely it was possible to record a complete TPP reversal in the absence of any SD events (**Fig. 2Aiii**). In 140 control preparations (53 males and 87 females) from 7 different datasets, within the 30 minutes cut-off, no SDs were recorded in 22 preparations (16%) and no TPP reversal, defined as ≥ 5 mV negative to the initial TPP, was recorded in 66 preparations (47%). After mechanically desheathing a ganglion using fine forceps (n = 4) the effect of ouabain was rapid and complete (**Fig. 2B**). This was not a routine procedure because of the likelihood of damaging the underlying neurons and neuropil. Ouabain-induced SDs were the same magnitude and had the same onset trajectory as SDs induced by anoxia (nitrogen suffocation) (**Fig. 2C**). However, whereas anoxic SD was accompanied by a complete cessation of neural activity recorded peripherally, in preparations that had not been rendered hypoxic by damaging the tracheae, neural activity and the ventilatory motor pattern could be recorded essentially unaltered during a ouabain SD event (**Fig. 2C**).

**Figure 2:**
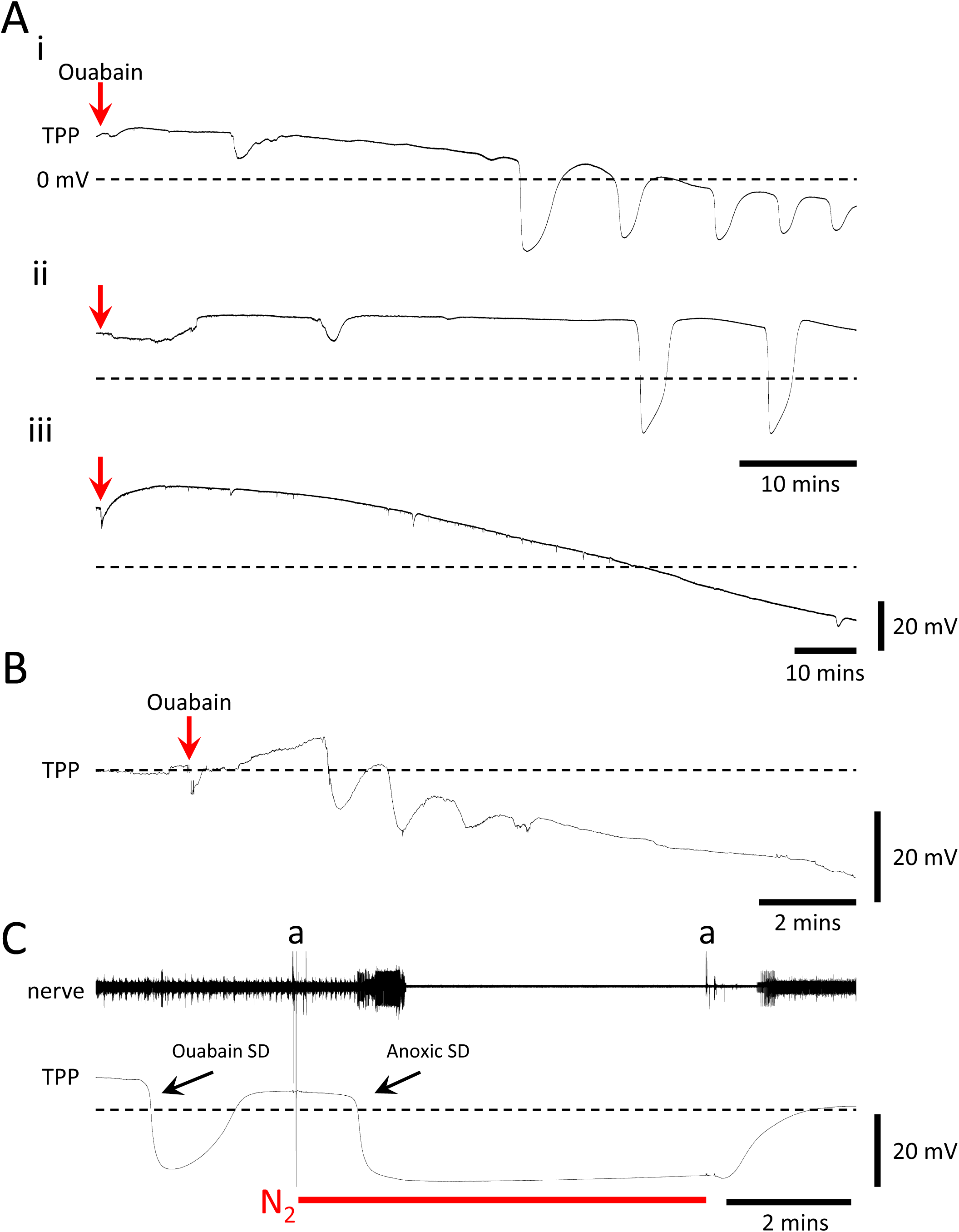
Phenomenology of ouabain-induced SD. Sample recordings showing: **Ai.** Usual TPP trajectory after bath application of 10^-2^ M ouabain (red arrow) with SD events occurring after a latency and superimposed on a gradual TPP reversal from positive to negative; **Aii.** SD events in the absence of TPP reversal; **Aiii.** TPP reversal in the absence of SD events (note the longer duration of the trace). **B.** Rapid TPP reversal after application of ouabain in a mechanically de-sheathed ganglion. **C.** Similarity between amplitudes and the onset and offset trajectories of ouabain-induced SD and anoxic SD induced by suffocation with N_2_ gas (timing indicated by red bar under trace). Note that anoxic SD affecting the whole ganglion reversibly shut off motor patterning and neural activity recorded extracellularly in a peripheral nerve, but this did not occur during the local ouabain-induced SD. Ouabain was administered before the start of the recording. Note the gradual TPP reversal in the background. **a** – electrical artifacts associated with turning on and off the flow of nitrogen.

These observations suggest that TPP reversal and repetitive SD are independent phenomena although both are induced by ouabain treatment. In addition, the ganglion sheath is effective at hindering ouabain’s access to the neuropil. Moreover, a ouabain-induced SD has a local effect and may not spread throughout the ganglion, for example to neuropil regions generating ventilatory motor activity. To investigate this further we undertook simultaneous, paired recordings from the same ganglion.

### Double penetration

Paired recordings of ouabain-induced SD were taken from the right and left sides of a ganglion in 12 preparations (3 male mesothoracic ganglion; 3 male metathoracic ganglion; 3 female mesothoracic ganglion; 3 female metathoracic ganglion). There is no reason to think that the uniformity of the simultaneous recordings from different locations will differ according to sex or ganglion identity. Whereas the initial TPP and minimum TPP (the trough of an SD event) were almost the same in right and left recordings, there was considerable variability in the trajectory of TPP reversal (from positive to negative) over the course of the recording (**Fig. 3**). Initial TPP was 16.2 ± 3.3 mV (mean ± SD; n = 24 recordings in 12 animals) and the mean absolute difference between simultaneous right and left recordings was 1.8 ± 1.8 mV. Similarly, the minimum TPP was −28.6 ± 4.5 mV and the mean difference between simultaneous right and left recordings was 4 ± 4.8 mV. On the other hand, TPP30 was −13.2 ± 17.9 mV with a mean difference between simultaneous right and left recordings of 20.0 ± 16.1 mV.

**Figure 3:**
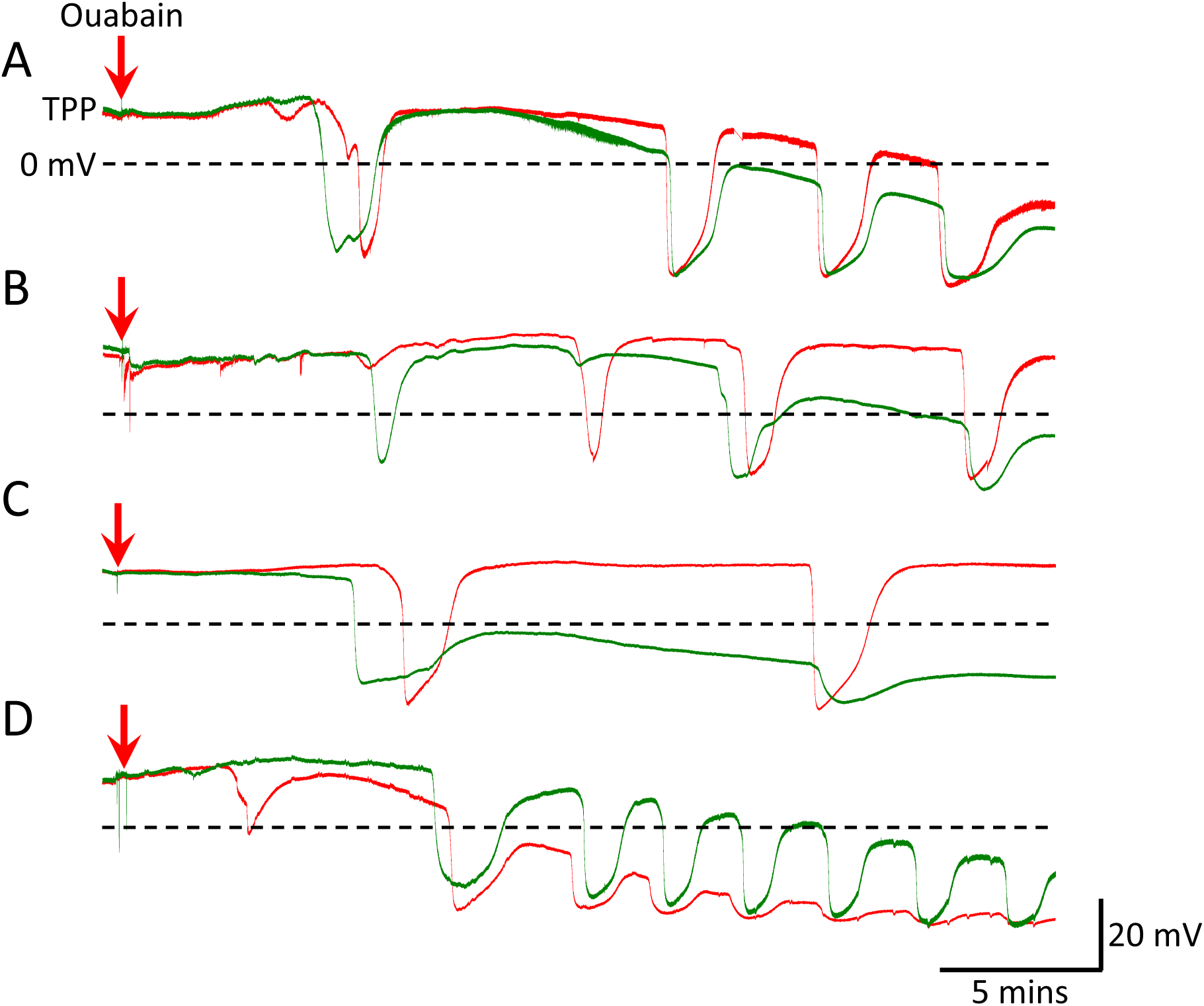
Parameters of TPP reversal and SD events varied independently. Overlays of sample recordings from different locations on the right and left sides of the same ganglion show that after bath application of 10^-2^ M ouabain (red arrows) the rate of TPP reversal was not obviously correlated with occurrence, timing, or amplitude of SD events. Note that the initial TPP after penetration was the same in both recordings and that there was a minimum TPP that determined amplitude of SD events. Left hemiganglion – red traces; Right hemiganglion – green traces.

Regarding the SD events themselves, the main difference between the right and left recordings was in the timing and occurrence of the first SD. In 4 of the 12 preparations the first SD was recorded in one electrode but not the other (see e.g. **Fig. 3B**). However, in all the preparations, after the initial discrepancy, an SD on one side occurred nearly simultaneously with an SD on the other side, indicating spread throughout the ganglion. This could be almost masked by the SD having a small amplitude due to occurring when TPP reversal was almost complete (see e.g. **Fig. 3D**). It is important to note that at no point during the slow TPP reversal did it ever recover and trend positive again. This is true for this dataset and for hundreds of preparations of ouabain-induced SD in many other datasets.

### TPP trajectory correlations

The results from double penetrations suggest that the trajectory of TPP reversal was more affected by the nature of the penetration than was the trajectory of SD events. During 30 minutes of ouabain treatment the gradual TPP reversal was associated with a decrease in the amplitude of SD events (**Fig. 4Ai**). However, scaling a small amplitude later SD to be the same amplitude of as a large amplitude early SD suggests that the timing of an SD was not affected by TPP reversal (**Fig. 4Aii**). We investigated the relationship between TPP reversal and SD parameters in a separate dataset of 45 middle-aged (∼5 weeks old) female locusts that had been collected previously.

**Figure 4:**
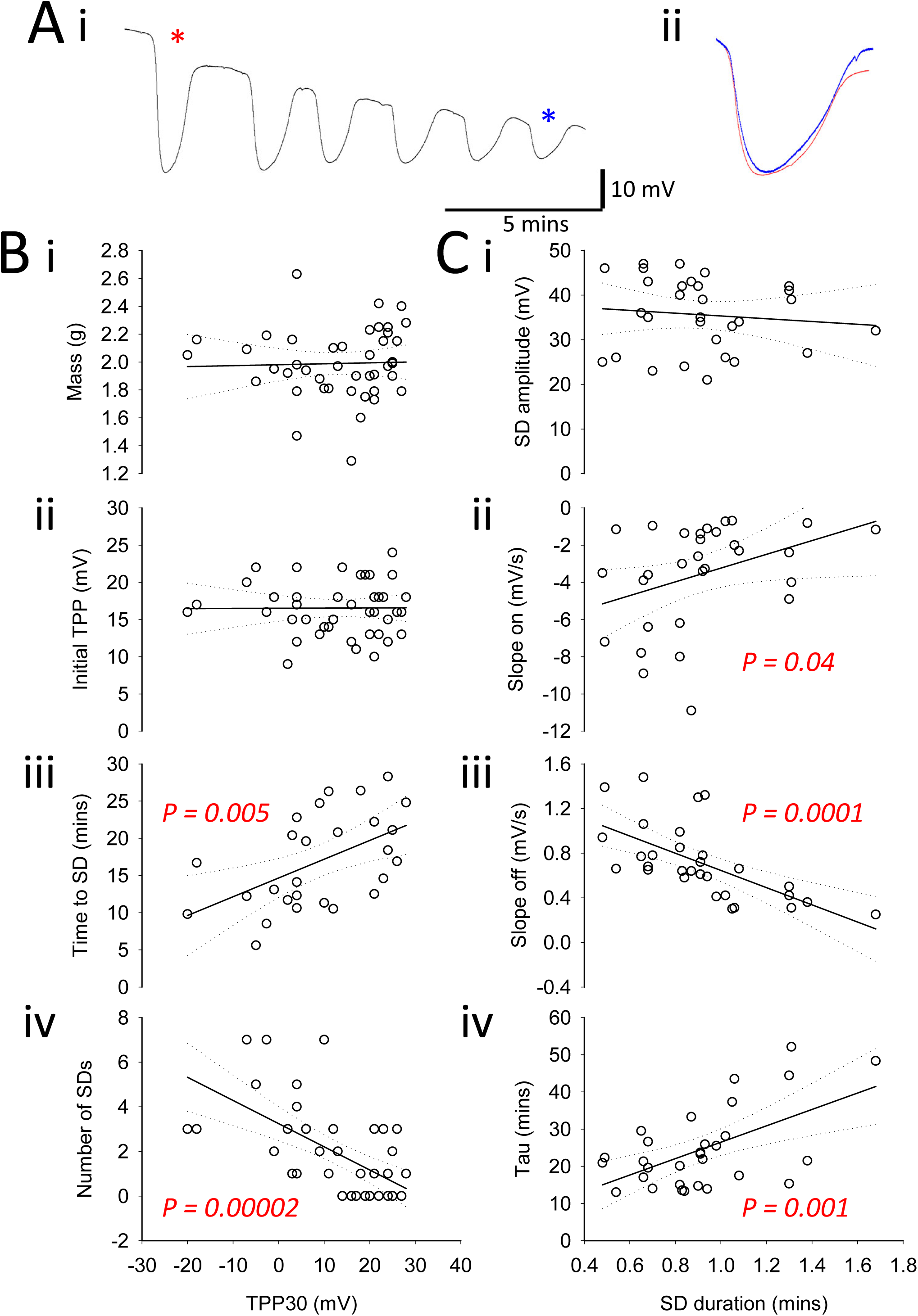
No correlation between rate of TPP reversal and timing of SD events. **A.** Sample recording of ouabain-induced repetitive SD. Individual events indicated by asterisks (red and blue) **(i)** are scaled to have equal amplitude and overlaid **(ii)**. Note that timing of the late, small amplitude event (blue) is identical to timing of the early, large amplitude event. Pearson correlations of data from female locusts: **B.** TPP30 is not correlated with mass **(i)** or the initial TPP **(ii)** but is positively correlated with the latency to the first SD event **(iii)** and negatively correlated with the number of SD events recorded in 30 minutes **(iv)**. **C.** SD duration is not correlated with SD amplitude **(i)** but is positively correlated with SD slope on **(ii)**, negatively correlated with SD slope off **(iii)** and positively correlated with tau, the time constant of the exponential recovery of TPP **(iv)**. Note that slope on is a negative value and thus increased duration was correlated with reduced rate of TPP change at both onset and offset of SD. Straight lines show linear regressions and dotted lines indicate 95% confidence limits. See also Table 1.

TPP reversal (reflected by TPP30) was not correlated with mass or initial TPP but was positively correlated with the latency to the first SD and negatively correlated with the number of SDs recorded in 30 minutes (**Fig. 4B**). Not surprisingly, the timing of SD events (reflected by SD duration) was correlated with measures of the rate of TPP change (slope on, slope off, and tau) but there was no correlation between SD duration and SD amplitude (**Fig. 4C**). Direct comparison between TPP reversal and SD parameters shows that there was no correlation between TPP30 and parameters of the SD trajectory although the measures of SD timing were significantly correlated with each other (**Table 1**).

**Table 1:** No correlation between rate of TPP reversal and timing of SD events. TPP30 was not correlated with slope on, duration, slope off, or recovery tau of SD events although these measures were significantly correlated with each other. R: Pearson correlation coefficient; P values in red indicate significant correlations.

|  |  | Slope on | Duration | Slope off | Tau |
| --- | --- | --- | --- | --- | --- |
| TPP30 | **R** | **0.006** | **-0.21** | **0.21** | **-0.064** |
|  | *P* | *0.98* | *0.28* | *0.18* | *0.69* |
| Slope on | **R** |  | **0.37** | **-0.4** | **0.008** |
|  | *P* |  | *0.044* | *0.031* | *0.97* |
| Duration | **R** |  |  | **-0.64** | **0.56** |
|  | *P* |  |  | *0.00014* | *0.0012* |
| Slope off | **R** |  |  |  | **-0.46** |
|  | *P* |  |  |  | *0.0015* |

The results described above indicate that TPP reversal and SDs are distinct phenomena although both are caused by ouabain treatment.

### Aging and SD

Extracellular K^+^ concentration (**Fig. 5A**) and TPP (**Fig. 5B**) recorded during ouabain-induced repetitive SD showed that aging slowed the recovery phase of SD but did not appear to affect the amplitude of events. We investigated this further in a dataset of 20 locusts (5 young males, 5 young females, 5 old males, 5 old females). One old male was removed as an outlier. Two-way ANOVAs revealed no effect of sex for any of the measures, so the graphs depict only the effects of age (**Fig. 6**).

**Figure 5:**
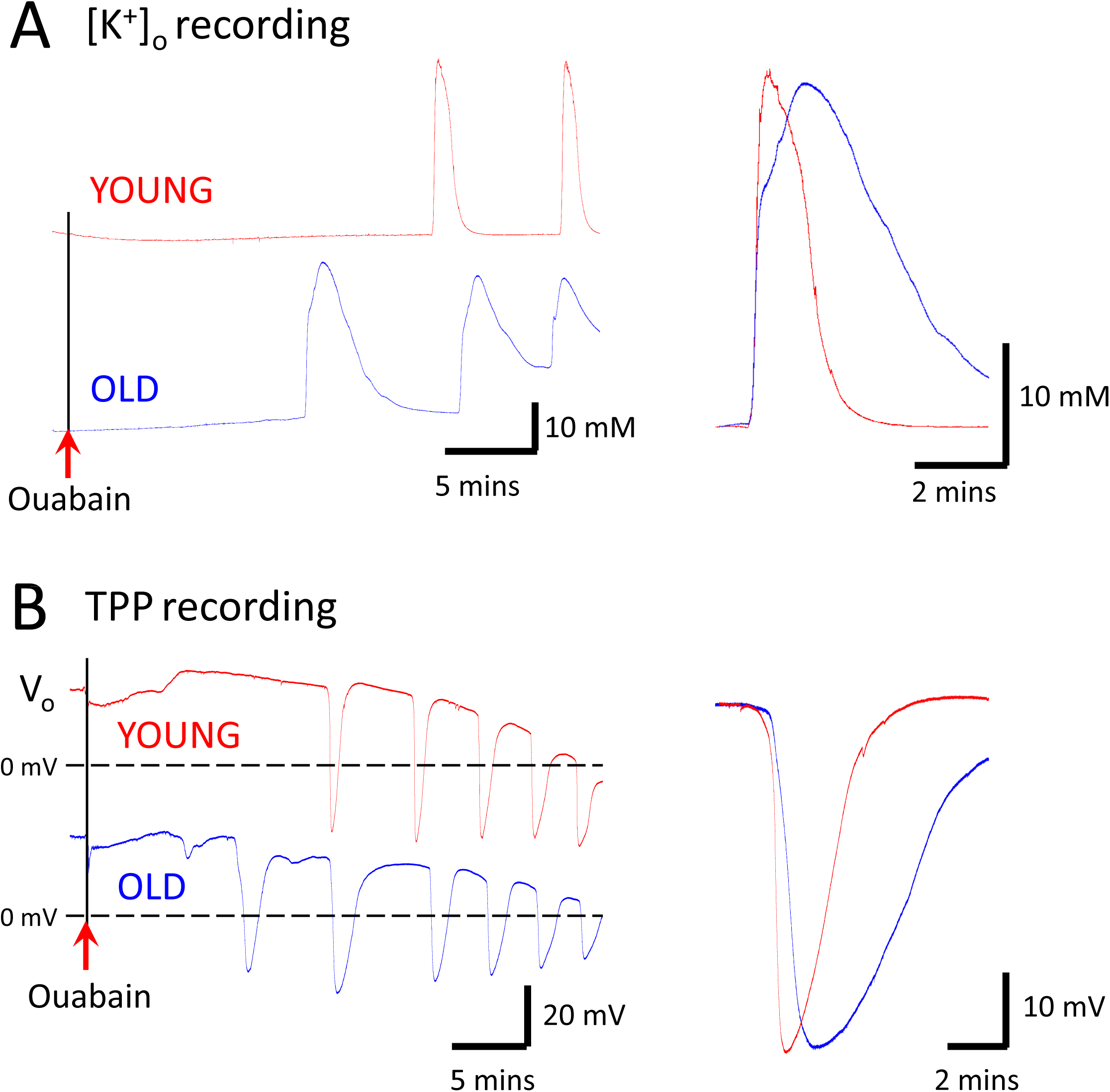
Ouabain induced repetitive SD in semi-intact preparations from young and old locusts. Sample recordings taken from the metathoracic ganglion in semi-intact preparations of young and old locusts show that bath application of 10^-2^ M ouabain induced **A.** repetitive surges of [K^+^]_o_ and **B.** abrupt and repetitive negative DC shifts of TPP (= extracellular potential relative to zero potential of the bath; V_o_), characteristic of SD. Overlays of traces at the same scale on the right shows that preparations from older locusts take longer to recover from SD disturbances of equal amplitude. Note also that V_o_ gradually reversed during the ouabain treatment. Young – red traces; Old – blue traces.

**Figure 6:**
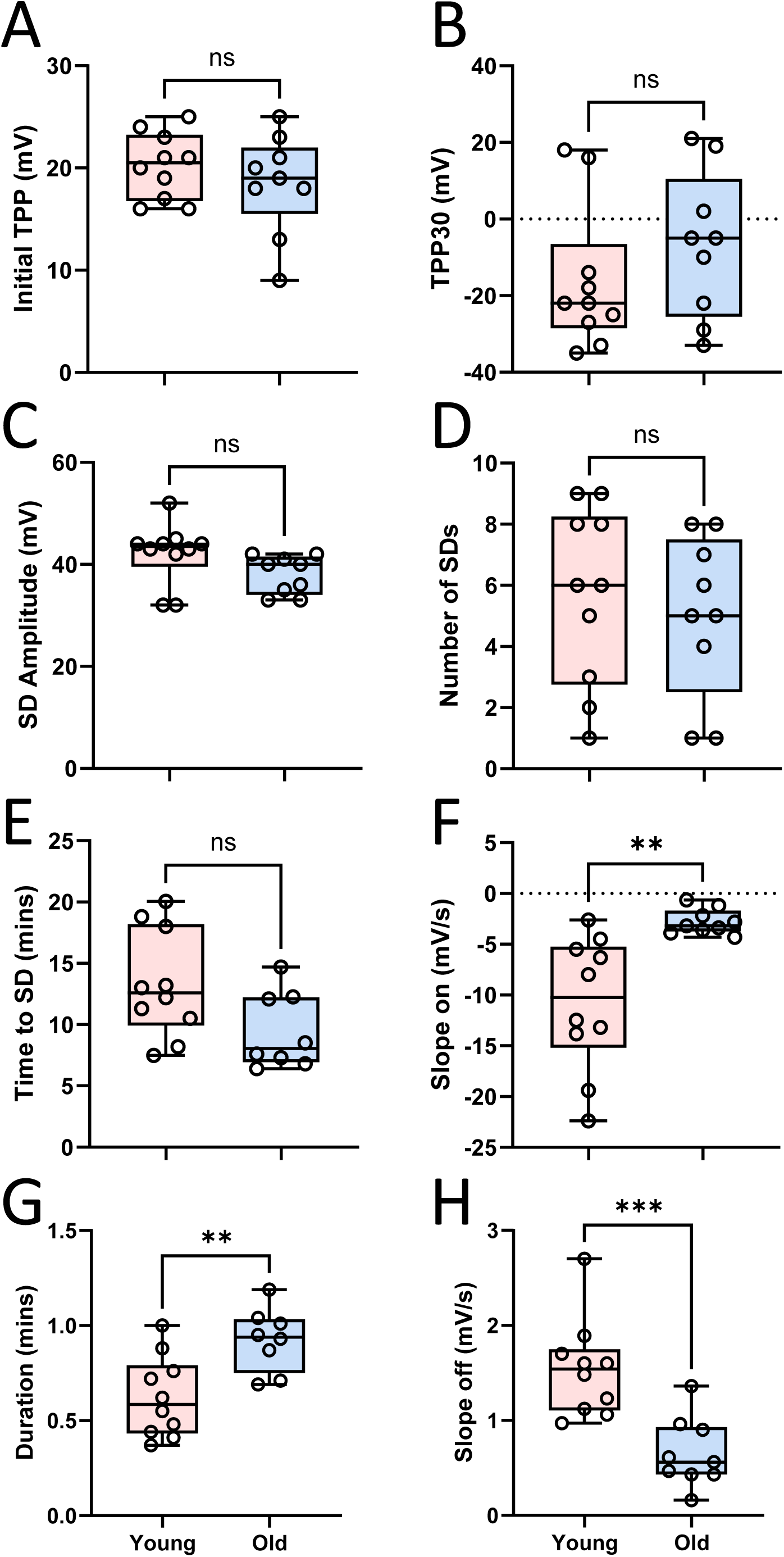
Aging slowed the timing of SD events induced by ouabain but did not affect the rate of TPP reversal. Data from male and female locusts combined: **A.** Initial TPP; **B.** TPP30; **C.** SD amplitude; **D.** number of SDs in 30 minutes; **E.** time to the first SD; **F.** SD slope on (note values are negative); **G.** SD duration; **H.** SD slope off. Statistics in text. Asterisks indicate significant differences using 2-way ANOVAs with no effect of sex and no interaction (*, P < 0.05; **, P < 0.01: ***, P < 0.001). Box plots indicate median, inter-quartile range, and whiskers to the minimum and maximum. Individual data points are shown as open circles.

Aging had no statistically significant effect on initial TPP, TPP30, amplitude of first SD, number of SDs in 30 minutes or latency to the first SD (**Fig. 6A-E**). SD in old locusts had a slower onset slope (mean ± SE; old = −2.8 ± 0.4 mV/s; young = −10.8 ± 2.1 mV/s; Welch’s t-test for unequal variances P = 0.004), a longer duration (old = 0.92 ± 0.06 min; young = 0.62 ± 0.07 min; Student’s t-test P = 0.005), and a slower recovery slope (old = 0.65 ± 0.4 mV/s; young = 1.54 ± 0.5 mV/s; Student’s t-test P = 0.0005) (**Fig. 6F-H**).

### Oxidative stress and SD

We investigated the effect of rotenone to increase oxidative stress in a dataset of 22 middle-aged locusts (5 control males, 5 control females, 6 rotenone males, 6 rotenone females). Initially we pretreated with 0.5 mM rotenone and increased to 1 mM rotenone midway through data collection. There was no significant effect of rotenone on TPP reversal or any of the SD parameters (**Table 2**).

**Table 2:** Bath applied rotenone (0.5-1.0 x 10^-3^ M) did not affect ouabain-induced SD. Two-way ANOVAs show no effect of sex or treatment on TPP reversal or timing of SD events. See text for details. SE: standard error.

|  | **Control (n=10)** |  | **Rotenone (n=12)** |  | 2-way P |
| --- | --- | --- | --- | --- | --- |
|  | mean | SE | mean | SE |  |
| **TPP (mV)** | 18.6 | 2.1 | 19.2 | 1.9 | 0.84 |
| **TPP 30 (mV)** | 6.6 | 6.1 | -3.8 | 5.5 | 0.22 |
| **SD amp (mV)** | 35.2 | 2.0 | 38.6 | 1.7 | 0.22 |
| **# SDs** | 3.4 | 0.8 | 4.3 | 0.7 | 0.44 |
| **time to SD (mins)** | 13.4 | 2.5 | 13.8 | 2.2 | 0.91 |
| **slope on (mV/s)** | -3.3 | 1.4 | -4.1 | 1.2 | 0.70 |
| **duration (mins)** | 0.8 | 0.1 | 0.9 | 0.1 | 0.30 |
| **slope off (mV/s)** | 1.0 | 0.1 | 0.8 | 0.1 | 0.43 |

On the other hand, pretreatment with an antioxidant, 2 mM NACA, in a dataset of 20 middle-aged male locusts (10 control, 10 NACA) did affect TPP reversal and recovery from SD (**Fig. 7**). There was no significant effect of NACA on initial TPP, SD amplitude, latency to the first SD and SD onset slope (**Fig. 7A, C, E & F**). NACA slowed TPP reversal, increasing TPP30 (**Fig. 7B**; mean ± SE; control = −17.3 ± 2.3 mV, NACA = 0.8 ± 5.2 mV; Student’s t-test P = 0.007) and this was associated with a reduction in the number of SDs in 30 mins (**Fig. 7D**; control = 4.6 ± 0.9, NACA = 2.3 ± 0.6; Student’s t-test P = 0.04). NACA also decreased the duration of the first SD (**Fig. 7G**; control = 1.07 ± 0.08 mins, NACA = 0.73 ± 0.06 mins; Student’s t-test P = 0.006) and increased the rate of SD recovery (**Fig. 7H**; control = 0.54 ± 0.10 mV/s, NACA = 1.0 ± 0.12 mV/s; Student’s t-test P = 0.016). It is worth noting that three NACA-treated locusts did not generate any SDs within the 30 minutes of ouabain treatment though all control locusts did.

**Figure 7:**
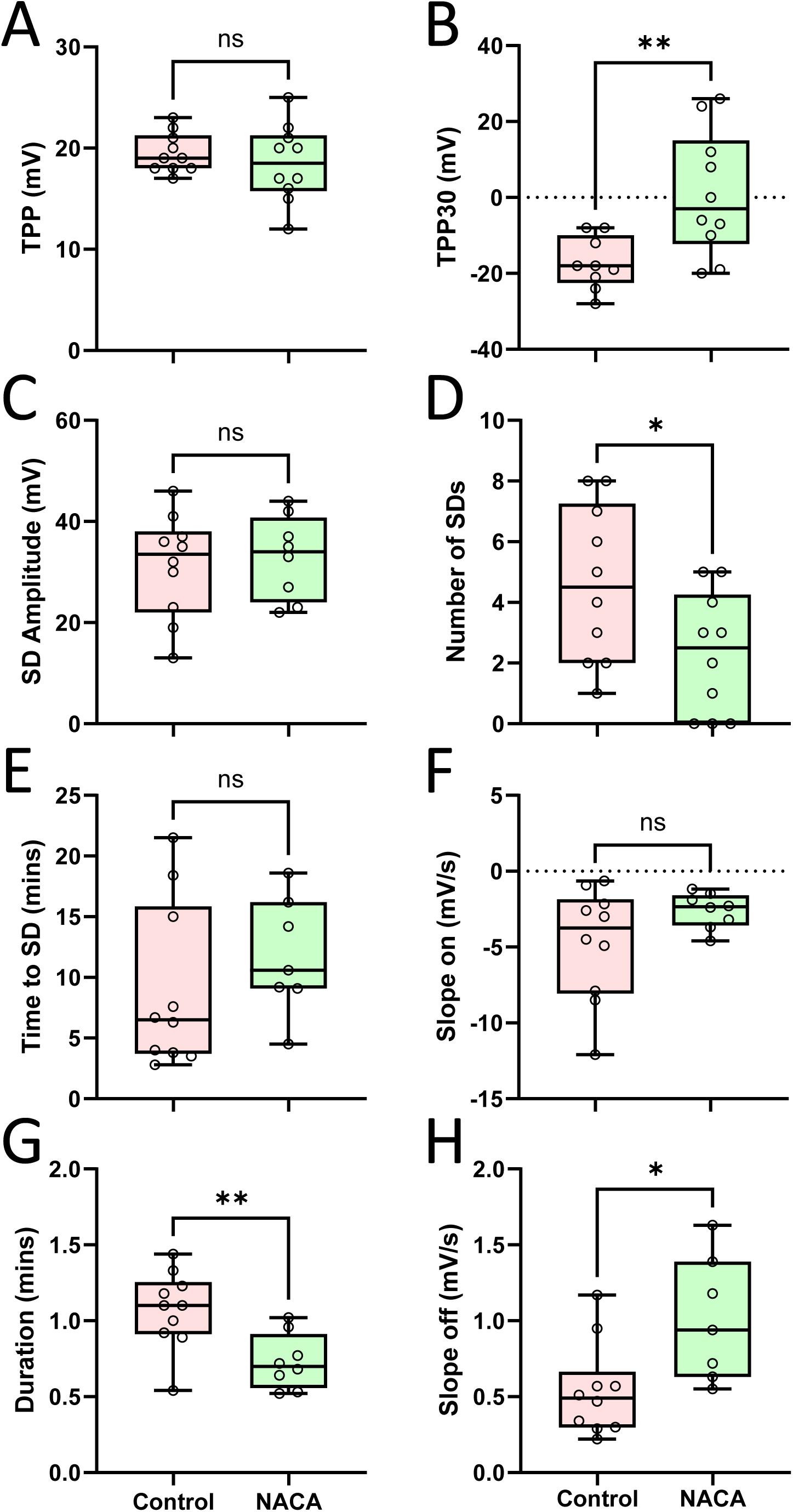
N-acetylcysteine amide (NACA, antioxidant) delayed TPP reversal and increased the rate of recovery of SD events. Data from male locusts: **A.** Initial TPP; **B.** TPP30; **C.** SD amplitude; **D.** number of SDs in 30 minutes; **E.** time to the first SD; **F.** SD slope on; **G.** SD duration; **H.** SD slope off. Statistics in text. Asterisks indicate significant differences using 2-way ANOVAs with no effect of sex and no interaction (*, P < 0.05; **, P < 0.01). Box plots indicate median, inter-quartile range, and whiskers to the minimum and maximum. Individual data points are shown as open circles.

## Discussion

Our main results can be summarized as follows. Aging impaired the ability of intact locusts to resist anoxia and to recovery from anoxia. Ouabain treatment of the CNS caused a gradual deterioration of ion homeostasis (TPP reversal) as well as abrupt SD events. These were independent of each other and likely had different underlying causes. Within the time constraints of our experimental procedure, aging did not affect the gradual deterioration but impaired SD recovery, i.e., the restoration of ion gradients and membrane potentials. Antioxidant treatment improved SD recovery, speeding the restoration of ion gradients, and delayed the gradual deterioration in the presence of ouabain.

We were interested in comparing young locusts with aged locusts, i.e. biological age rather than chronological age. Aging can be characterized in terms of lifespan (percent survival), which in locusts is associated with the accumulation of neurolipofuscin arising from oxidative damage of mitochondria (Fonseca et al., 2005). On a 14:10 (L:D) photoperiod, gregarious *Locusta migratoria* have a maximum lifespan of ∼33 days post-imaginal moult with abnormal mitochondria evident in muscle at 28 days (Guo et al., 2021). With this photoperiod, young locusts can be characterized as 14 days (at 80% survival) and old locusts as 28 days post imaginal moult (at 10% survival), associated with increased mitochondrial ROS and accelerated muscle aging (Guo et al., 2023). This gives a 2-week difference between young and old locusts. However, lifespan is strongly affected by photoperiod, and lifespan is longer under a 12:12 photoperiod compared with a 16:8 (L:D) photoperiod (Rodgers et al., 2006). Gregarious, male *Schistocerca gregaria* on a 12:12 (L:D) photoperiod have been designated as old at 24 days compared with young at 10 days, which is just after completion of maturation (Austin et al., 2024). This also gives a 2-week difference between young and old locusts, and, based on the percent survival of locusts and median life expectancy of humans, the age of the old locusts was cautiously estimated to be equivalent to a human age of ∼73 years (Blockley et al., 2022). In our aging experiments, old locusts were >3 weeks older than young ones; we are confident that our old locusts were aged.

The mass of intact locusts affected their ability to resist submersion anoxia and to recover when returned to normoxia. Young locusts showed a negative correlation between time to coma and mass. The time to coma in adult locusts is positively correlated with tracheal volume (Rodgers-Garlick et al., 2011) and negatively correlated with body mass (Robertson et al., 2019). Within an instar, tracheal volume decreases with increasing mass (Lease et al., 2006; Robertson et al., 2019), attributable to an age-related gain in mass within a restricting cuticle. Hence, the lightest in the “young locusts” group were likely the youngest with a relatively large tracheal volume and consequently an improved “breath-holding” ability. Indeed, late stage (∼4 weeks post imaginal moult) mature *Schistocerca americana* are more vulnerable to hypoxia and have reduced aerobic capacity compared to early stage mature locusts (Greenlee and Harrison, 2004). Old locusts showed a negative relationship between measures of recovery and mass. An explanation for this is that the lightest in the “old locusts” group were likely the oldest that had experienced more deleterious effects of aging such as muscle wastage, reducing mass. Thus, there was variability in the extent of aging within the cohort of old locusts and the oldest of these were impaired in their ability to recover from anoxia compared with young locusts.

Although SD is the basis for the anoxic coma, it is important to remember that recovery from anoxia in intact animals is a progression of recoveries that occur sequentially, of: ion homeostasis, neural membrane potentials, neuron excitability, synaptic transmission, motor coordination and muscle function (Robertson and Van Dusen, 2021). The timing of SD is relevant primarily for only the first two of these and measurements of TPP (onset and recovery) are marginally relevant for the whole animal responses to anoxia, although recovery of TPP is the *sine qua non* of the progression.

In insects the TPP is measured as the voltage difference across the ion diffusion barrier of the sheath, between the haemolymph (or bathing solution) and the interstitium of the CNS. It results from the difference between the basolateral membrane potential (facing the haemolymph) and the adglial membrane potential (facing the neuropil) (Schofield and Treherne, 1984; Robertson et al., 2020). Thus, TPP changes can reflect changes in either or both basolateral and adglial membrane potentials and these, in turn, reflect changes in ion conductances and ion concentration gradients across them. The abrupt large (∼40 mV) negative shift of the TPP is coincident with a surge of [K^+^]_o_ and is interpreted as reflecting a collapse of ion gradients in the neuropil and depolarization of the adglial membrane potential to ∼0 mV. During ouabain-induced SD, the rapid TPP transients usually ride on top of a gradual negative shift of TPP which reflects a simultaneous gradual increase of [K^+^]_o_. This is also true for repetitive SD induced by fluorocitrate in rat brain (Largo et al., 1997). However, we found that SD and TPP reversal could be recorded independently of each other, although it was rare to record complete TPP reversal without SD. The fact that mechanical de-sheathing of the ganglion resulted in a rapid, complete TPP reversal suggests that the integrity of the sheath, and thus ouabain’s ease of entry to the neuropil, influenced the rate of TPP reversal. Paired TPP recordings showed little correlation between TPP reversal at two sites on either side of the same ganglion although, once started, SD events were evident simultaneously at both electrodes. The simplest explanation for the difference in TPP reversal at the two sites is that the quality of the electrode penetration was different, with penetration damage being associated with more rapid TPP reversal. Moreover, analysis of a large dataset showed no correlation between the rate of TPP reversal and any timing parameters of SD event, though there were strong correlations between SD timing parameters themselves. It is a pertinent observation that there was never any recovery from TPP reversal, no matter how little it had progressed. In this respect it resembles the negative ultraslow potential (NUP), which can be recorded from mammalian brains, including those of terminal human patients. The NUP is defined as: “*A very long-lasting, shallow negativity of the DC potential with superimposed SDs. Experimentally associated with incomplete recovery of the typical ion changes after SDs and hence with developing neuronal injury. NUP may indicate that only a fraction of neurons in the tissue have repolarized at the recording site and that the remaining fraction is persistently depolarized*.” (Dreier et al., 2017). Moreover, it has been suggested that the large size of the NUP in humans could reflect a drift of the DC potential across the blood brain barrier due to tissue damage or cell death (Dreier et al., 2019) and the TPP is measured across the locust blood brain barrier. Previously we have suggested that the gradual effect of ouabain on the TPP indicates a gradual inhibition of NKA as ouabain diffuses through the ganglion (Van Dusen et al., 2020). The current results show that TPP reversal is locally generated and stabilizes at the trough of SD events. Given our observations and the similarities with the NUP, we propose that TPP reversal in our experiments indicates a time- and concentration-dependent, local and permanent impairment of ion homeostasis by ouabain, which can be exacerbated by damage caused by electrode penetration, with some penetrations being more damaging than others.

Perhaps not surprisingly, the TPP onset and offset trajectories of ouabain-induced and anoxia-induced SD were similar but, whereas nerve activity, generated centrally but recorded peripherally, was terminated during anoxic SD it was unaffected during ouabain SD. Previous recordings of ouabain-induced SD have described shutdown of nerve activity (Rodgers et al., 2007; Rodgers-Garlick et al., 2011) however, for historical reasons these recordings were made in preparations that had damage to the tracheae supplying the metathoracic ganglion (by raising the ganglion on a metal plate) probably rendering it hypoxic. This is evident in the different SD event durations, which were ∼1 min in the current experiments (without the plate). In previous experiments SD duration was ∼2 mins, moreover compound C (an inhibitor of the energy sensor, AMPK) reduced duration, which is associated with motor pattern recording during SD events (Rodgers-Garlick et al., 2011). We suggest that in a relatively undamaged, non-hypoxic, ganglion, short duration SD events, particularly at the start of SD generation, may not propagate throughout the ganglion thus sparing some circuitry from shutdown.

The discussion above leads to the conclusion that ouabain causes two separable phenomena: a slow permanent negativity (NUP) and abrupt reversible negativities (SD). Within the time course of our experiments, aging affected only the latter of these, lending support to conclusion that their underlying causes are different. Older animals had slower and longer duration SD events but no change in TPP reversal. Such an increase in duration and reduction in the rate of DC potential recovery of SD (KCl-induced) has been recorded after ischemia in the rat brain and this is potentiated by aging (Menyhárt et al., 2015). However, aging did not change these measures in non-ischemic rat brains.

Our experiments with agents to induce or relieve oxidative stress were limited by the fact that we did not deliberately investigate interactions between age and oxidative stress. For example, it would have been informative to examine the effects of rotenone on young animals and the effects of NACA on old animals in a controlled manner. Retrospectively comparing the chronological age of locusts (days since post imaginal moult) in the aging dataset with those in the rotenone and NACA datasets (Kruskal-Wallis One Way ANOVA P < 0.0001), rotenone ages (median 32.5 days) were different (Dunn’s P < 0.001) from the old group (median 51 days) but not (Dunn’s P = 0.12) from the young group (median 26 days), whereas the NACA ages (median 38 days) were different (Dunn’s P < 0.001) from the young group but not (Dunn’s P = 0.08) from the old group (**Fig. 8**). This analysis should be treated cautiously because the datasets were collected separately and there may have been colony differences. Nevertheless, it seems that the rotenone experiment used younger locusts and the NACA experiment used older locusts.

**Figure 8:**
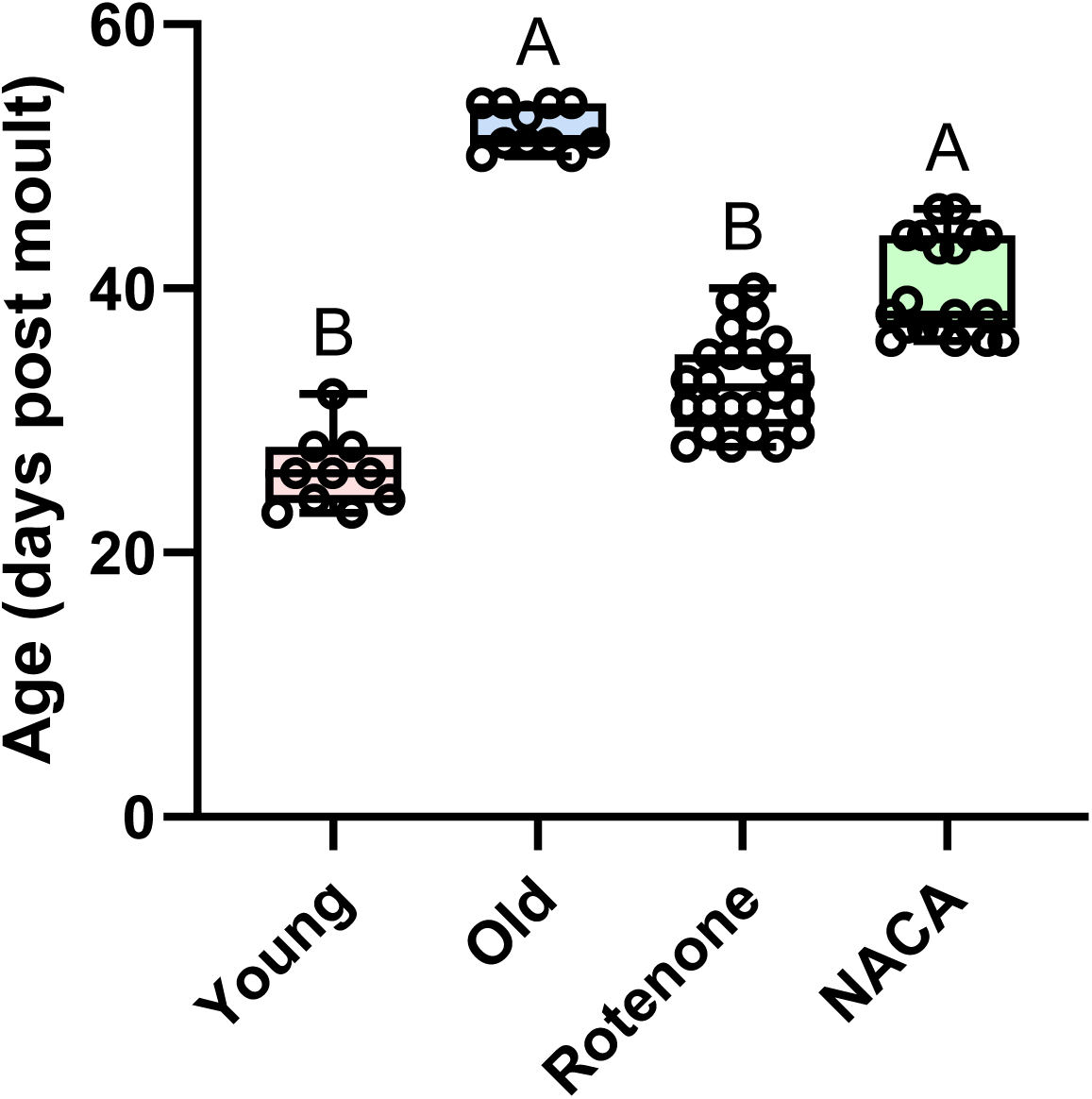
Age of locusts in datasets of experiments using semi-intact preparations. Box plots indicate median, inter-quartile range, and whiskers to the minimum and maximum. Individual data points are shown as open circles. Significant differences indicated by different letters (Kruskal-Wallis One-Way ANOVA followed by Dunn’s pairwise tests).

We were surprised that rotenone did not exacerbate SD, indeed had no observable effect, in spite of the fact that its ability to generate oxidative stress is well established (Areiza-Mazo et al., 2018) and it is a potent insecticide used to model Parkinson’s disease in animal and insect models (Richardson et al., 2019). Moreover, rotenone is known to increase the propagation speed of cortical spreading depression in rat brains (Amaral de Brito et al., 2020) and SD in mouse brain slices (Germano et al., 2022), although aging generally decreases SD propagation speeds (Guedes and Abadie-Guedes, 2019). We suspect that the lack of an effect of rotenone in our preparation was due to the concentration we used and predict that more diligent examination with a range of concentrations would yield positive results.

Treatment with an antioxidant had the opposite effect to aging, decreasing SD duration and increasing the slope of TPP recovery, suggesting that NACA prevented damage during ouabain treatment and that this effect of aging was due to oxidative stress. The antioxidant treatment also slowed TPP reversal and decreased the number of SDs recorded during 30 minutes of ouabain. One possibility is that the mechanisms that cause TPP reversal, which, being permanent, is a more consequential outcome, are less vulnerable to aging but still responsive to antioxidant. There is an abundance of literature relating aging, oxidative stress and brain function in mammals, including using cortical spreading depression and SD to monitor effects of antioxidants and aging (Guedes et al., 2012; Guedes and Abadie-Guedes, 2019). Nevertheless, there have been relatively few studies that focus on SD recovery, particularly with reference to cellular mechanisms. The current consensus in mammals is that the effect of aging on SD recovery is associated with oxidative damage to NKA and there is a suggestion that glial clearance mechanisms may be involved (Hertelendy et al., 2019). There is little doubt that any disorder of NKA will have a profound effect on restoration of ion gradients. Properties of this important enzyme are regulated by many different mechanisms (Moyes et al., 2021) and modifications to NKA by thermal acclimation change the temperature sensitivity of ouabain-induced SD, reflecting a change in the environmental chill tolerance (coma SD temperature) of *Drosophila* (Andersen et al., 2022). Regarding glial clearance mechanisms, the blockage of gap junctions in the locust CNS induces rapid repetitive SD ([K^+^]_o_ surges) associated with a gradual increase in background [K^+^]_o_ and also exacerbates the effect of ouabain (Spong and Robertson, 2013). If aging affects glial clearance mechanisms via gap junctions it would certainly have consequences for ouabain-induced SD. Interestingly, the plateau of [K^+^]_o_ reached after blockage of gap junctions is not an end point. Addition of sodium azide to induce chemical anoxia causes a further increase to a second plateau of [K^+^]_o_ (Spong et al., 2014) and this may reflect NKA shutdown.

The similarity in the effect of aging on SD duration and recovery slope in locusts and mammals reinforces the belief that insects provide good models for rapid and invasive investigation of mechanisms of SD. Our results suggest that future research on the role of the blood brain barrier will be fruitful.

## Acknowledgements

We thank Hanna Grover for collecting the data in Figure 1, and Mads K Andersen and Heath MacMillan for their comments on a previous version of the manuscript. Funded by a Discovery Grant from the Natural Sciences and Engineering Research Council of Canada.

## Abbreviations

CNS: central nervous system
NACA: N-acetylcysteine amide
NKA: Na^+^/K^+^-ATPase
NUP: negative ultraslow potential
ROS: reactive oxygen species
SD: spreading depolarization
TPP: transperineurial potential
TPP30: transperineurial potential after 30 minutes of ouabain treatment

